# *Escherichia coli* Bcteriuria in pregnant women in Ghana: Antibiotic resistance pattern, Virulence Factors and Resistant genetic markers

**DOI:** 10.1101/317065

**Authors:** Forson Obeng Akua, Wilson Bright, David Nana-Adjei, Marjorie Ntiwaa Quarchie, Noah Obeng-Nkuramah

**Affiliations:** Department of Medical Laboratory Science, School of Biomedical and Allied Health Sciences, College of Health Sciences, University of Ghana, Legon, Accra, Ghana.

**Keywords:** *Escherichia coli*, bacteriuria, pregnant women, Ghana

## Abstract

The relevance of *Escherichia coli* associated bacteriuria infection in pregnant women is poorly understood, despite these strains sharing a similar virulence profile with other extra intestinal pathogenic *E. coli* producing severe obstetric and neonatal infections. We characterized and determined the antimicrobial susceptibility, resistant genes and virulence profiles of 82 *E. coli* isolates associated with asymptomatic bacteriuria in some pregnant in five very distinct hospitals in the Volta region from January, 2016 to April, 2016 using Kirby-Bauer disc diffusion and polymerase chain reaction.

High levels of antimicrobial resistance was observed to Ampicillin (79.3%), Tetracycline (70.7%) and Cotrimoxazole (59.8%), except for Cefuroxime (32.9%). Resistant genes analyses revealed 58.5% were positive for *Bla*_TEM_ and 14.6% for *aph(3)-Ia(*aphA2). Virulence factors (VFs) was more widespread in pregnant women in the 2^nd^ and 3^rd^ trimesters than 1^st^ trimester. VFs relating to adhesion (*pap*C and *iha*), Protectins (*tra*T), aerobactin acquisition (*iut*A) and iron acquisition systems (*fyu*A and *irp*2) were more prevalent in the resistant *E. coli* isolates. This study provides additional evidence for a link in bacteriuria and transmission of extra-intestinal *E. coli* in pregnant women to cause multi-resistant severe obstetric or neonatal infections. Considering the involvement of extra-intestinal *E. coli* in infections, our results may be helpful to develop strategies to prevent maternal and/ neonatal infections. In addition continuous surveillance is required to guide appropriate antibiotic usage in pregnant women.

## Introduction

Maternal genitourinary infection is a leading cause of pregnancy complications worldwide [1]. In the last decades, the rod shaped gram-negative lactose fermenter and gas producing member of the family *Enterobacteriaceae* called *Escherichia coli* is reported to be a major cause of UTI in pregnancy [2, 3]. There are many pathotypes of *E. coli*, but the pathotype associated with extra intestinal infection is called extra intestinal pathogenic *Escherichia coli* (ExPEC) [4, 5, 6]. ExPEC are characterized by the presence of large numbers of specialized virulent factors (VFs) that enable them to become invasive, adhesive, and resistant to bactericidal drugs or resistant to phagocytosis in the host [7, 8]. ExPEC lack the ability to cause gastroenteritis in humans [7, 9], however, they are able to cause extra-intestinal infections involving sepsis, meningitis, cellulitis, osteomyelitis, wounds infections and urinary tract [9, 10, 11].

ExPEC infections are common causes of healthcare-associated infection in recent years with a large number reported to be associated with bacteriuria, bacteraemia and urosepsis [12-15]. Urinary tracts infections (UTI) is treatable, however, it is becoming increasingly difficult to control because of rampant antimicrobial resistance to pregnancy friendly antibiotics, especially those belonging to the beta lactam class, cephalosporins and fluoroquinolones [13, 16, 17]. Although the prevalence of antimicrobial resistance in *E. coli* in pregnant women has been found to vary in India, Iraq, and Ethiopia [18, 19, 20, 21]. In Ghana, some hospitals have reported *E. coli* as one of the pathogens associated with bacteriuria and bacteraemia [22, 23, 24]. However, characterization of virulence factors and genetic properties of the associated pathogenic isolates are limited to basic phenotypic tests leaving several important questions unanswered on the implicated *E. coli* strain (s) infection and its propensity to cause other extra intestinal infection in patients. Hence the need for this study to evaluate the antibiotic resistance phenotypes, and virulence factor genes of ExPEC isolates from pregnant women with asymptomatic bacteriuria from randomly selected hospitals in the Volta region of Ghana.

## Materials and methods

### Study location

The Volta region is one of the administrative regions in Ghana with Ho as its capital. It lies east of the Volta Lake and about 20570 km^2^ [25]. The Volta regional hospital, Ho serves as a referral center for the Volta region in Ghana and also to some West African countries. There is an infection control unit in the hospital which supervises and coordinates hygienic practices to prevent and control outbreaks of multi-resistant pathogenic infections. The Volta region is bordered to the east by the Republic of Togo, to the south by the Atlantic Ocean and to the north by the Northern region of Ghana. Ghana has a population of about 24.2 million and is considered a lower middle income economy [30]. Health facilities available in the bacteriology laboratory of regional hospitals in Ghana permit only phenotypic characterization of bacteria and the most common organisms reported in the laboratories are *Escherichia coli*, *Staphylococcus aureus* and *Pseudomonas aeruginosa* [26]. *Escherichia coli* isolated in the Bacteriology Laboratory of the hospitals are not routinely tested for virulence factors or sequenced to detect the sequence type complex [27].

### Subject selection and data collection

Pregnant women attending antenatal clinic at the selected hospitals were only included in the study. Consenting women were then given written detailed information about the study and a written informed consent form to complete. A self-administered questionnaire was given to the women to obtain information on the demographic and socio-economic characteristics (see S1 File for Copy of Questionnaire). All information about the study was translated verbally in the native languages for those who could not read. Participants who could not write were also assisted to fill the consent forms. Non pregnant and pregnant women who were on antibiotic treatment were excluded from the study. The sample size was calculated using the formula; N=Z^2^P (1-P)/ D^2^ where; N= sample size; Z= 1.96 (95% confidence interval), P = 56.5%, D= 0.05 T

### Microbiological analysis

Clean catch midstream urine samples from the pregnant women were inoculated onto cysteine lactose electrolyte deficient (CLED) agar and incubated at 37°C for 18 to 24 hours. Colonies that appeared circular and yellow on CLED agar were considered to be potential *E. coli* [28]. A representative colony on each plate was Gram stained and further tested using indole, methyl red, citrate, Voges-Proskauer test and urease [28]. API 20E identification system (bioMerieux SA, Marcy l‟Etoile, France) was used to confirm the isolates before the identified isolates were stored in 10% glycerol-trypticase soy broth at −70°C for further sensitivity and molecular analyses testing.

### Antimicrobial susceptibility testing

The antimicrobial susceptibility testing was carried out on the isolates using the Kirby Bauer method based on the CLSI, (29) guidelines for resistance to Ampicillin (10μg), Tetracycline (30μg), Cotrimoxazole (25μg), Nalidixic acid (30μg), Nitrofurantoin (300μg, Gentamicin (10μg) and Cefuroxime (30μg).

The control strains used for the determination of minimum inhibitory concentrations were *E. faecalis* ATCC 29212, *E. coli* ATCC 25922, *Pseudomonas aeruginosa* ATCC 27853 and *Staphylococcus aureus* ATCC 29213. The bacteria were subcultured onto 5% horse blood agar (HBA) plates (37°C, 18 h) and then suspended in saline to a concentration equivalent to 0.5 Mcfarland. A loopful of the suspensions was transferred to a Mueller-Hinton agar plate and a sterile cotton swab was used to streak the entire surface of the plate. The lid of the agar plate was left ajar for 3 - 5 min to allow for excess surface moisture to be absorbed before the application of drug impregnated disks and incubated at 24°C for 24 hr. After incubation, zone diameters around the antibiotic discs was measured and classified as sensitive or resistant based on the CLSI [29] break points (Table 1).

**Table 1.** Guidelines for interpreting antimicrobial susceptibility results

|  | Zone diameter (mm) |  |
| --- | --- | --- |
|  | Susceptible | Resistant |
| Ampicillin (10) | $\geq 17$ | $\leq 13$ |
| Tetracycline (30) | $\geq 15$ | $\leq 11$ |
| Cotrimoxazole (25) | $\geq 16$ | $\leq 10$ |
| Nalidixic acid (30) | $\geq 19$ | $\leq 13$ |
| Nitrofurantoin (300) | $\geq 17$ | $\leq 14$ |
| Gentamycin (10) | $\geq 15$ | $\leq 12$ |
| Cefuroxime (3) | $\geq 23$ | $\leq 14$ |

### DNA extraction

A single colony of a fresh bacterial culture from 5% HBA was picked and suspended in 200 ml of sterile water. Tubes were heated at 98°C for 10 min and subsequently centrifuged at 17 900 X g for 5 min. The supernatant were recovered and 2 ml of this was used as a template in the various polymerase chain reactions (PCR).

### Molecular Analysis of *E. coli* isolates

A total of 82 *E. coli* isolates recovered from the pregnant women from the five hospitals in the Volta region of Ghana between February, 2016 to August, 2016 were analysed for virulence factors (Vfs) with PCR for the presence of genes encoding 18 VFs (30, 31). The following genes: adhesins (*pap*C, *pap*G, including *pap*G alleles, *sfa*/*foc*, *iha*, *hra* and *ibe*A), toxins (*hlyC*, *cnf1* and *sat*), iron capture systems (*fyuA, irp2, iroN, iucC and ireA*), protectins (*neu*C, chromosomal *ompT* and *traT*) and *usp*, a gene encoding uropathogenic-specific protein were tested using primers (Integrated DNA Technologies, Inc, USA - https://www.idtdna.com) in Table 2.

**Table 2.**
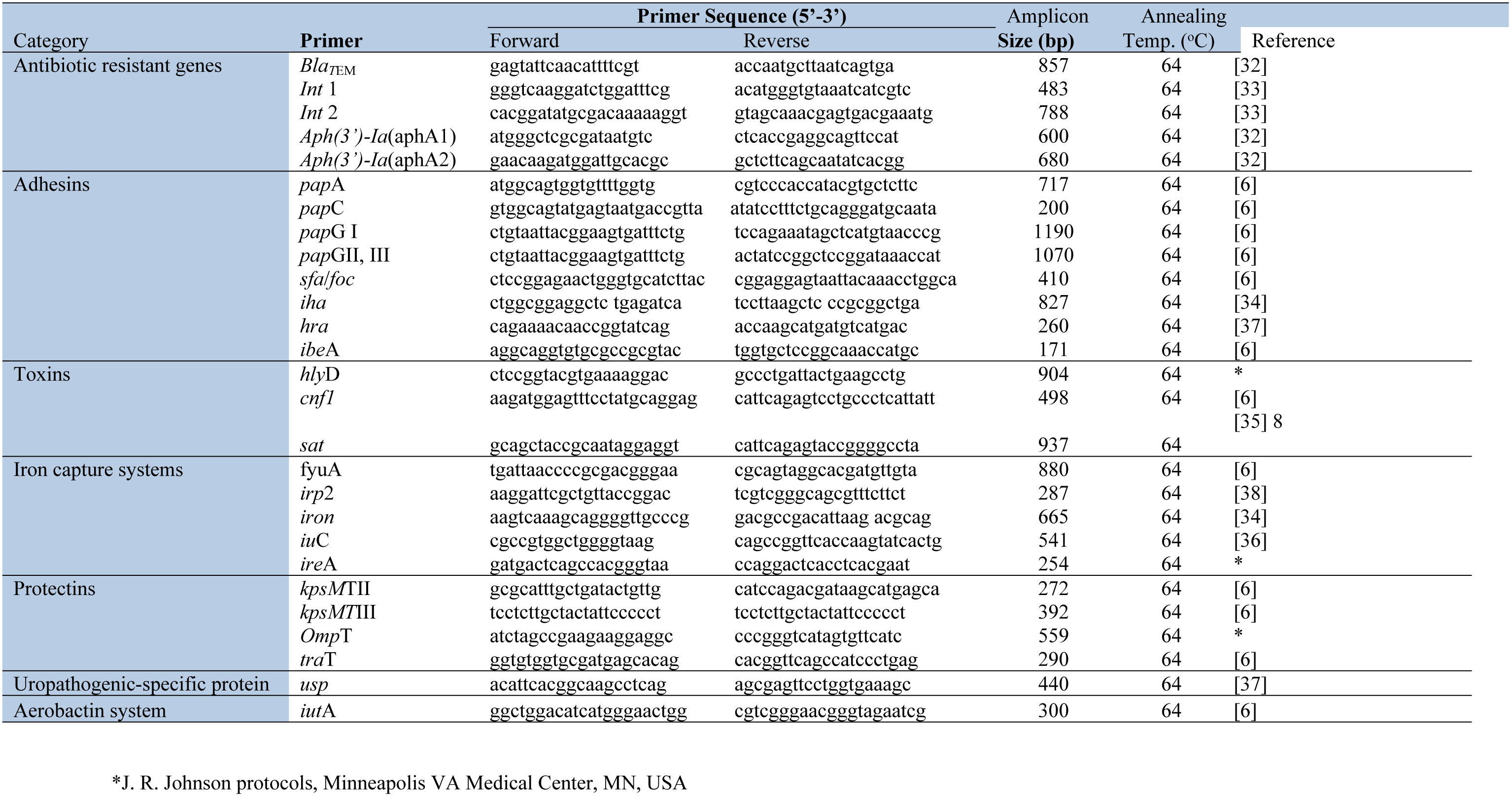
Primers used for PCR.

Each reaction consisted of 4 mM MgCl_2_, 1 ml of 25 pmol of each primer, 2 ml of 2 mM dNTPs and 4 ml of 5 X PCR buffer, 1U of Taq DNA polymerase (New England BioLab, South Africa) in a total reaction volume of 25 ml, including 2 ml DNA template. Six primer pools utilised were; pool 1: *iron* (665), *sfa* (410), *iut*A (300), *hra* (260), pool 2: *pap*A (717), *KpsM*TIII (392), *ire*A (254), *ibe*A (171); pool 3: *pap*G1 (1190), *pap*GII, III (1070), *iha* (827), *omp*T (559), *KpsM*TII (272); pool4: *iuC* (541), *Cnf*1 (498), *irp*2 (287); pool 5: *hly*D (904), *usp* (440), *tra*T (290); and pool 6 : *pap*C (200), *sat* (937), *Fyu*A (880).

The cycling condition were as follows: 94 °C for 5 min, followed by 30 cycles of denaturation (94 °C, 30 s), annealing (64°C, 30 s), extension (68°C, 3 min) and final extension (72°C, 10 min). PCR products were then electrophoresed on 1.5% agarose gel containing ethidium bromide.

### Antibiotic resistant genes determination

Resistant genes for the various phenotypic resistant strains were determined using primers and corresponding annealing temperatures highlighted in Table 2. Each reaction mixture consisted of 4 mM MgCl_2_, 1 ml of 25 pmol of each primer, 2 ml of 2 mM dNTPs and 4 ml of 5 X PCR reaction buffer, 1U of Taq DNA polymerase (New England BioLab, South Africa) in a total reaction volume of 25 ml, including 2 ml DNA template. DNA amplification was carried out using the following conditions: 7 min initial denaturation at 95 °C, following 35 cycles of denaturation at 94 °C for 30 s, annealing at various temperatures for 30 s (Table 2) and extension at 72 °C for 45 s. PCR products were then electrophoresised on a 1.5% agarose gel containing ethidium bromide. The size of the various amplicons was determined by comparison with 100 bp and 1 kb ladders.

### Data handling and statistical analysis

The data were entered into Microsoft Excel and analyzed using GraphPad Prism software, version 6. In all cases, P-values less than 0.05 were considered statistically significant. Initially the association between each exposure and the presence of infection was assessed using the Chi-squared test. Chi-square analysis was carried out to test for significance between prevalence of intestinal parasitic infections and risk factors for prevalence of intestinal parasitic infections. Odds ratios were computed to measure the strength of association. To determine independent risk factors for infection, logistic regression analysis was employed where appropriate.

### Ethical approval

The study was approved by the Ethics Committee of the School of Biomedical and Allied Health Sciences, College of Health Sciences, University of Ghana, Legon (Ethics Identification Number: SAHS/10507884/AA/MLS/2015–2016). Participation was voluntary and written consent was taken in accordance with the ethical committee’s guidelines. Permission was also sought from the Volta Region Ghana Health Service, all participating Hospitals and laboratory personnel before the samples were taken.

## 3.0 RESULTS

### 3.1 Distribution of UTI and socio-demographic characteristics

Out of the 400 urine specimens from pregnant women in the five selected hospitals, 42.8% (171) of the pregnant women were positive for bacteriuria (growth >10^5^ colony forming units/mL), whilst 57.25% (229) had no significant growth. *E. coli* formed majority of the microbes associated with bacteriuria (48%), but low prevalence were found for *Staphylococcus aureus* (18.1%), *Klebsiella pneumoniae* (13.45%), *Proteus mirabillis* (11.11%), *Pseudomonas aeruginosa* (5.26%), and *Enterococcus faecalis* (4.09%) (Table 3).

**Table 3.**
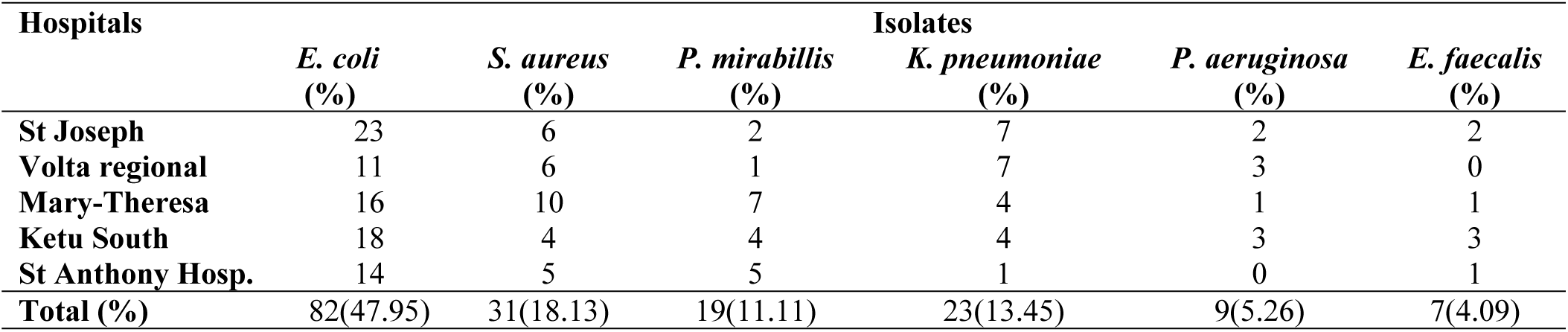
Distribution of isolated bacteria in the different hospitals

The educational levels of the four hundred pregnant women revealed basic level education (62.7%), and secondary education (25.5) were the common levels of education (Table 4). One hundred and ten (43.8%) of the women with basic education had UTI and 45.5% (50) were infected with *E. coli*. Forty five percent of the pregnant women with secondary education had UTI and *E. coli* was associated with 52.2% (24). Only 15 (31.9%) women with tertiary education had UTI and 8 (53.3%) of the cases of UTI was caused by *E. coli*. Despite the differences in the rate of UTI in relation to educational levels of the participant, the difference was non-significant (χ^2^ (2, N = 400) = 2.678, p = 0.262).

**Table 4.** Socio-demographic characteristics and distribution of UTI among pregnant women

| Characteristics | No. of Pregnant Women Tested (%) | Culture results | | | $\chi^2$ | p value |
| --- | --- | --- | --- | --- | --- | --- |
|  |  | No of Significant growth (%) | No. of no significant growth | No. of UTI associated with *E. coli* (%) |  |  |
| **AGE (years)** |  |  |  |  |  |  |
| 13 - 19 | 54 (13.5) | 24 (44.4) | 30 (55.6) | 8 (33.3) | 1.397 | 0.706 |
| 20 - 29 | 228 (57.0) | 96 (42.1) | 132 (57.9) | 50 (52.1) |  |  |
| 30 - 39 | 102 (25.5) | 42 (41.2) | 60 (58.8) | 21 (50.0) |  |  |
| 40 - 49 | 16 (4.0) | 9 (56.2) | 7 (43.8) | 3 (33.3) |  |  |
| **GESTATIONAL AGE** |  |  |  |  |  |  |
| 1<sup>st</sup> Trimester | 71 (17.8) | 20 (28.2) | 51 (71.8) | 11 (55.0) | 12.209 | 0.002* |
| 2<sup>nd</sup> Trimester | 156 (39.0) | 66 (42.3) | 90 (57.7) | 35 (53.0) |  |  |
| 3<sup>rd</sup> Trimester | 173 (43.3) | 85 (49.1) | 88 (50.9) | 36 (42.4) |  |  |
| **PARITY** |  |  |  |  |  |  |
| Nulliparous | 129 (32.3) | 52 (40.3) | 77 (59.7) | 30 (57.7) | 1.025 | 0.599 |
| Primiparous | 107 (26.8) | 44 (41.1) | 63 (58.9) | 18 (40.9) |  |  |
| Multiparous | 164 (41) | 75 (45.7) | 89 (54.3) | 34 (45.3) |  |  |
| **EDUCATION** |  |  |  |  |  |  |
| Basic Level | 251 (62.7) | 110 (43.8) | 141 (56.2) | 50 (45.5) | 2.678 | 0.262 |
| Secondary | 102 (25.5) | 46 (45.1) | 56 (54.9) | 24 (52.2) |  |  |
| Tertiary | 47 (11.75) | 15 (31.9) | 32 (68.1) | 8 (53.3) |  |  |
| **Hospitals** |  |  |  |  |  |  |
| St Joseph Hospital, Nkwanta | 80 (20.0) | 42 (52.5) | 38 (47.5) | 23 (54.76) | 9.847 | 0.043* |
| Volta Regional Hospital, Ho | 80 (20.0) | 28 (35.0) | 52 (65.0) | 11 (39.29) |  |  |
| Mary Theresa Catholic Hospital, DodiPapase | 80 (20.0) | 39 (48.7) | 41(51.3) | 16 (41.03) |  |  |
| Ketu South Municipal Hospital, Aflao | 80 (20.0) | 36 (45.0) | 44 (55.0) | 18 (50.00) |  |  |
| St Anthony Hospital, Dzodze | 80 (20.0) | 26 (32.5) | 54 (67.5) | 14 (53.84) |  |  |
| **Total** | **400 (100.0)** | **171 (42.8)** | **229 (57.2)** | **82 (47.95)** |  |  |

\*Significant at <0.005

For the purpose of this study, the participants were put into 4 age groups (Table 4). Occurrence of UTI among 13 - 19 age groups was found to be 44.44% and 33.33% of these UTIs were associated with *E. coli*. Out of the 228 patients (aged 20 - 29 years), 96 (42.11%) of the pregnant women were found to have UTI with 50 (52.08%) associated with *E. coli*. The rate of UTI in the 30 - 39 age groups was 42.18% and *E. coli* was associated with 50% (20 pregnant women). The age group 40 – 49 years had 9 participants (56.25%) positive bacteria growth with *E. coli* causing 33.33% of the UTI (Table 3). The differences in the rate of UTI among the various age groups was statistically non-significant (X2 (3, N = 400) =1.397, p = 0.706).

Most of the pregnant women were in their third [43.3% (173)], and second [39% (156)] trimesters of pregnancy (Table 4). Significant bacteria growth of 42.3% (66) and 49.1% (85) was recorded for pregnant women in their second and third trimesters. Fifty-five per cent (11) of the significant bacteria growth recorded among first trimester group was for *E. coli*. Other cases of UTI caused by *E. coli* were found to be (53.0% (35) and 42.3% (36) respectively for second and third trimesters. Chi square exact test performed to determine the relationship between the gestational age and the development of UTI revealed a significant association with gestational age (X2 (2, N = 400) = 12.209, p = 0.002).

The distribution of parity of participants was 32.3% (129), 26.8% (107), and 41% (164) for nulliparous, primiparous and multiparous respectively. *E. coli* was isolated from 22.2%, 66.7% and 38.5% of urine samples collected from nulliparous, primiparous and multiparous pregnant women respectively.

### 3.2 Antimicrobial susceptibility pattern of *E. coli* isolates

The 82 *E. coli* isolates from the pregnant women revealed high resistance to ampicillin (Table 5). A resistant prevalence of 86.96%, 72.7%, 68.8%, 88.2% and 78.6% to *E. coli* isolates from St Joseph hospital, Volta regional hospital, Mary Theresa Hospital, Ketu south Municipal hospital and St Anthony hospital respectively was found. *E. coli* however recorded the least resistance of 26.09%, 27.27%, 18.75% and 21.43% to Cefuroxime for isolates collected from St Joseph hospital, Volta regional hospital, Mary Theresa Hospital and St Anthony hospital respectively. A relatively high resistance of 70.59% for cefuroxime to *E. coli* isolates was observed in Ketu South hospital, Aflao (Table 5).

**Table 5.** Antibiotic resistance pattern of *E. coli* from five hospitals in the Volta region, Ghana.

| ANTIBIOTIC | HOSPITALS (No.) |  |  |  |  |  |
| --- | --- | --- | --- | --- | --- | --- |
|  | St Joseph hospital<br>(n=23, %) | Volta regional hospital<br>(n=11, %) | Mary Theresa hospital<br>(n=16, %) | Ketu south Mun. Hospital<br>(n=18, %) | St Anthony hospital<br>(n=14, %) | Total<br>(n=82, %) |
| Ampicillin | 20 (86.96) | 8 (72.73) | 11 (68.75) | 15 (88.24) | 11 (78.57) | 65 (79.3) |
| Tetracycline | 18 (78.26) | 8 (72.73) | 10 (62.50) | 12 (70.59) | 10 (71.43) | 58 (70.7) |
| Cotrimoxazole | 16 (69.57) | 6 (54.55) | 8 (50.00) | 9 (52.94) | 10 (71.43) | 49 (59.8) |
| Nalidixic Acid | 8 (34.78) | 6 (54.55) | 8 (50.00) | 11 (64.71) | 7 (50.00) | 40 (48.8) |
| Nitrofurantoin | 7 (30.43) | 4 (36.36) | 5 (31.25) | 9 (52.94) | 4 (28.57) | 29 (35.4) |
| Gentamicin | 6 (26.09) | 4 (36.36) | 6 (37.5) | 10 (58.82) | 8 (57.14) | 34 (41.5) |
| Cefuroxime | 6 (26.09) | 3 (27.27) | 3 (18.75) | 12 (70.59) | 3 (21.43) | 27 (32.9) |

Resistance of 78.26%, 72.73%, 62.50%, 70.59% and 71.41% were recorded against Tetracycline for *E. coli* isolates from patients attending St Joseph hospital, Volta regional hospital, Mary Theresa Hospital, Ketu south Municipal hospital and St Anthony hospital (Table 5). Similarly, at the St Joseph hospital, 69.57%, 34.78%, 30.43%, and 26.09% of *E. coli* isolates were resistance to Cotrimoxazole, Nalidixic acid, Nitrofurantoin and Gentamicin respectively. Whilst at Volta regional hospital, 54.55% of the *E. coli* isolates were resistant to Cotrimoxazole and Nalidixic acid. Resistance to Nitrofurantoin and Gentamicin was found to be 36.36%.

### 3.4 Prevalence of Antibiotic Resistant Genes and integrons in the *E. coli* isolates

In total, forty seven ampicillin resistant isolates were found to contain *Bla*_TEM_ (Table 6). Pregnant women in the 2^nd^ (24 isolates) and 3^rd^ (18 isolates) trimesters had *E. coli* isolates with more *Bla*_TEM_ gene compared to women in their 1^st^ trimesters (5 isolates). The aminoglycoside genes *aph(3)- Ia*(aphA2) for gentamicin resistance was found in only 6 phenotypically resistant isolates from 6 pregnant women in their 2^nd^ and 3^rd^ Trimesters (Table 6). All the *E. coli* isolates were screened for the presence of *int*I and *int*II, however only 10 of isolates were positive for *int*I, whilst 2 E. coli isolates contained *int*II, 58 of the isolates did not possess either *int*I or *int*II.

### 3.5 Distribution of Virulence factors gene

The distribution of the 82 *E. coli* isolates in relation to virulence genes from the various groups of pregnant women revealed 75.6% (62 isolates) *E. coli* contained two or more virulence genes (VFs) (Table 6). The virulence score used to classify the ExPEC isolates was calculated using the total number of VFs genes. Isolates were classified as ExPEC if they were positive for two or more of the tested virulence genes (5). The *iut*A (aerobactin acquisition), *pap*C and *iha* (adhesins), *fyu*A and *irp*2 (iron capture systems), *tra*T (protectins) were the common detected genes, whereas *usp* (uropathogenic-specific proteins) and some of the adhesin genes (*hra*, *ibe*A, & *pap*G1) were the least detected genes, *sat* (toxins) and *pap*GII & *pap*GIII (adhesion) was not detected in any of the isolates. In addition, VFs was more widespread in pregnant women in the 2^nd^ (30 isolates) and 3^rd^ (25 isolates) trimesters than 1^st^ trimester (12 isolates).

**Table 6.**
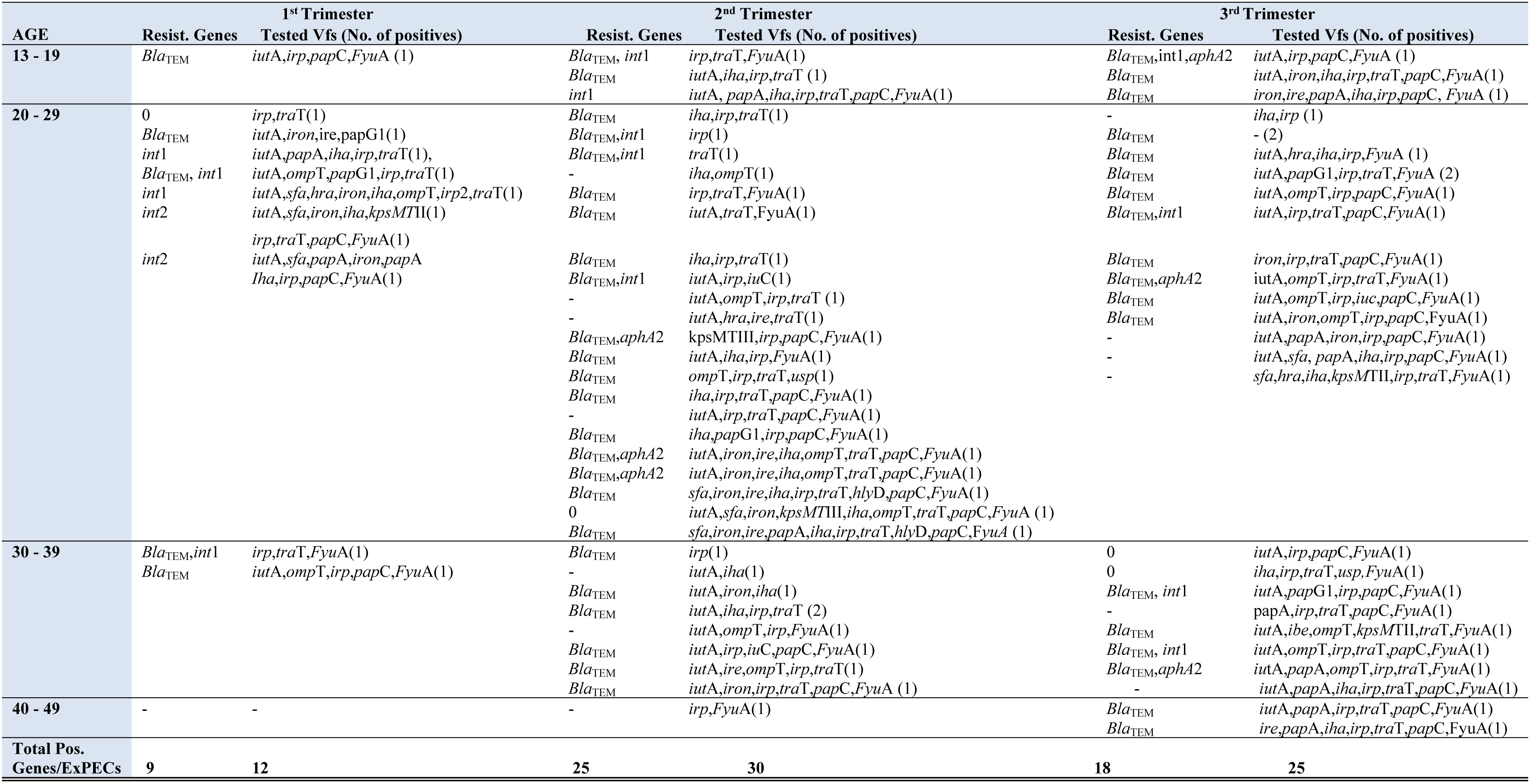
Distribution of Virulence factors (VFs) and resistant genes in *E. coli* isolates among the different trimester of pregnant women.

In the tested pregnant women, all the *E. coli* isolates in pregnant women in the 1^st^ Trimester were ExPEC isolates. Although pregnant women aged between 20 - 29 years were more positive compared to women aged between 13-19 years and 30 -39 years, pregnant women in their 2^nd^ trimesters aged between 20 -29 years (21 patients) had a higher prevalence of VFs. Out the 33 VFs positive *E. coli* isolates associated with pregnant women in their 2^nd^ trimester, 30 of the strains were positive for two or more of the tested virulence genes. Whilst a few (2) pregnant women aged 40 - 49 in their 3^rd^ trimesters haboured ExPEC isolates, 12 of the women aged between 20 -29 were positive for ExPEC isolates.

## 4.0 DISCUSSION

There are few studies on the antimicrobial susceptibility and/or virulence of *E. coli* isolates colonizing the genital tract of pregnant women [39, 40, 41]. However, no studies have been carried out to systematically compare virulence factors and antimicrobial resistance in *E. coli* isolates from pregnant women in different trimesters in Ghana.

This study revealed 42.75% of the 400 sampled pregnant women had UTI. This prevalence (42.75%) is slightly lower than the 56.5% prevalence reported in a previous study in Ghana in Cape Coast by Alex *et al*., [42] in 200 asymptomatic pregnant women. Although finding in this study are quite similar to 47.5% by Okonko *et al*., [43] in Nigeria, it is lower than the 85% prevalence reported in Edo state, Nigeria by Turay *et al*., [44]. However, the prevalence of 42.75 reported in this study is higher compared to the 7.3% reported in Kumasi by Turpin *et al*., [45], 5.1% by Lumbiganon *et al*., [46] in Thailand and 18.8% prevalence of UTI in Southern Ethiopia [47]. The difference in the prevalence in the different countries can be attributed to varied genital hygiene and socioeconomic conditions [48].

*E. coli* accounted for 47.95% of the UTI cases recorded in this study. This is in conformity with previous studies by Hamdan *et al.*, [49] in Sudan, Kawser *et al.*, [50] in Bangladesh, Akobi *et al*., [51] in Nigeria and Alex *et al.*, [42] in Ghana that *E. coli* is the major cause of UTI in pregnancy. The high incidence of *E. coli* associated with UTI among the pregnant women can be attributed to poor genital hygiene practices that enable movement of *E. coli* from its natural habitats into the genitals [52].

The predominance of *Staphylococcus aureus* (18.13%) over *Klebsiella pneumoniae* (13.45%) in this study is in contrast to Alex *et al.*, [42] study which reported *Klebsiella pneumoniae* as the second prevalent among asymptomatic pregnant women. This result can be attributed to varying social, biological and environmental factors which facilitate host system diversity in different countries [52]. The high numbers of *Staphylococcus aureus* in this study may suggest that *Staphylococcus aureus* is gaining clinical grounds as one of the common etiological agent of UTI in pregnancy and further studies are required to validate the its potential to cause infectious diseases in pregnancy.

### 4.1 Socio-demographic characteristics and UTI among pregnant women

Many socio-demographic factors are known to affect the frequency of bacteriuria during pregnancy [53]. These factors can include multiparity, gestational age, previous medical history of UTI, diabetes mellitus and anatomic urinary tract abnormalities [53, 54].

This study revealed that participants in 40 – 49 age group had the highest UTI prevalence of 56.25% whilst the least incidence was 41.18% in 30 – 39 age group. Although statistical analysis revealed no significant differences in age groups and occurrence of UTI, findings from this study are in contrast to Sheerin [52] study which reported UTI development to be directly proportional to the age of the participants. Other underlying factors like personal hygiene and differences in levels of sexual activity among the various age groups may have affected the propensity to develop UTI in the tested pregnant women [52].

Pregnant women in their third trimester in this study recorded the highest incidence of UTI (49.13%), followed by those in their second trimester (43.25%). Finding are conformity with Kawser *et al*., [50], Ferede *et al*., [56], Al-Haddad [57], and Sibi *et al*., [58] studies from Bangladesh, Ethiopia, Yemen, and India which reported that prevalence of UTI occurs with increase in gestational age. Although Chi square exact test analysis revealed a statistical association (*p* = 0.002) of prevalence of UTI and gestational age in this study. Findings however are in contrast to studies by Turay *et al*., [44] and Onuh *et al.*, [21] from Nigeria and Alex *et al*., [42] in Ghana which reported a higher prevalence of UTI in the second trimester compared to the third trimester.

The prevalence of UTI was found to increase with parity in this study, though the difference observed in this study was found to be non-significant (p = 0.599). Multiparous pregnant women had the highest incidence of UTI with 45.12%, and this was followed by 42.06% and 40.31% for primiparous and nulliparous pregnant women respectively. This finding however, are in contrast to findings by Emiru *et al*., [53] and Nandy *et al*., [59] that increase in parity is directly proportion to increase in susceptibility to UTI.

Although a chi square exact test revealed no significant association between educational level and development of UTI in pregnancy (*p* = 0.262), pregnant women with secondary education (45.1%) and basic level of education (43.8%) were positive for bacteriuria. The changes in the rate of UTI among the participants in various level of education may be as result of varying levels of health education on personal hygiene practices; however our findings are similar to Emiru *et al*., [53] study in Ethiopia.

### 4.2 Phenotypic antimicrobial resistance and resistant genes

In this study, *E. coli* isolates were found to have high antimicrobial resistance against Ampicillin, Tetracyclines, and Cotrimoxazole. These prevalences are similar to earlier report by Newman *et al*., [22], Feglo [60] and Newman *et al.*, [60] in Ghana. This high level of resistance observed in this study can be attributed to abuse of these drugs over the years. This is because they are relatively cheap and easily accessible without prescription [22, 61]. Although most of the *E. coli* isolates were susceptible to cefuroxime (33.33%). This finding however contradict Okonko *et al*., [43] study which reported higher resistance of *E. coli* to Cephalosporins than Quinolone/ Fluoroquinolones. The difference in the findings can be attributed to differences in geographical locations with varying levels of exposure to antibiotics. Although no significant differences was recorded between the antimicrobial resistances from the selected facilities in the Volta region, the difference in resistance pattern of the *E. coli* isolates in the different hospitals may be due to environmental factors, societal factors including and indiscriminate use of antibiotics among the general populace [22)]

### 4.3 Prevalence of Virulence factors genes and antibiotic resistant genes

We recognised a considerable number of the bacteria harboured the *iut*A (aerobactin acquisition), *pap*C and *iha* (adhesins), *fyu*A and *irp*2 (iron capture systems), *tra*T genes [62, 63]. In contrast to Sáez-López *et al*., [41] study with pregnant women, the ExPEC isolates in this study showed high antimicrobial resistance as previously reported in studies in some African countries [64, 65]. In addition, the ampicillin-resistant ExPEC isolates containing *Bla*_TEM_ gene showed a greater number of VFs in comparison with tetracycline or gentamicin resistant isolates, being highly significant for *iut*A, *irp*, *tra*T, and *ih*A. Whilst our findings are similar to Sáez-López *et al*., [41] study which evaluated *E. coli* colonizing the vagina and causing obstetric infection in pregnant women in Barcelona, it is dissimilar to Ramos *et al*., [66] study with *E. coli* causing UTI in pregnant women in Sweden, Uganda, and Vietnam. The differences in the studies may be due to varying geographical area, host physiological changes or susceptibility to *E. coli* isolates with pathogenic islands containing VFs.

## 5.0 Conclusion

In conclusion, our results demonstrate that the ability to adhere, invade and utilize the iron acquisition systems are important for antibiotic resistant ExPEC isolates that are associated with UTI in Ghanaian pregnant women. Whilst treating UTI infections in pregnant women in low income countries can be challenging due increasing resistance to first line drugs, extensive use and misuse of antibiotics [22], the appropriate empiric treatment and clinical management of pregnant women is mandatory to provide the appropriate interventions to avoid the aetiological link between maternal symptomatic or asymptomatic carriage of pathogens with obstetric infections. Considering the involvement of extra-intestinal *E. coli* in infections, our results may be helpful to develop strategies to prevent maternal and/ neonatal infections.

### Limitations

One limitation of the present study is that it focused on asymptomatic infection rather symptomatic infection and included only five hospitals from different locations of a single Region in Ghana, thereby not allowing extrapolation of our results to other Regions in country or other African countries. However, this is the first study to characterize VFs gene and resistant genes in *E. coli* isolates from pregnant women in the country. Furthermore, we were unable to follow up the pregnant patients to determine outcome of treatment of asymptomatic infection on maternal and neonatal health.

## Acknowledgments

The authors would like to express their gratitude to all the staff of St Joseph, Volta regional, Mary-Theresa, Ketu South and St Anthony Hospitals and all the pregnant women for their cooperation and support during the various aspects of the study. Special thanks to Dr. James R. Johnson, Adam L. Stell and Brian Johnston of the University of Minnesota, Department of Medicine and Infectious Diseases for providing the positive controls for the virulence factors.

## Availability of data and materials section

The datasets used and/or analyzed during the current study are available from the corresponding author on reasonable request.

## Funding

This research received no specific grant from any funding agency in the public, commercial, or not-for-profit sectors.

### Consent for publication

Not applicable

## References

1. Gilbert N. M., O’Brien V. P., Scott Hultgren, Macones G., Lewis W. G., Lewis A. L. (2013). Urinary Tract Infection as a Preventable Cause of Pregnancy Complications: Opportunities, Challenges, and a Global Call to Action. Global Advances in Health and Medicine, 2 (5):59–69.

2. Ayub M., Amir J. S., Firdous K., et al. (2016). E. coli the Most Prevalent Causative Agent Urinary Tract Infection in Pregnancy: Comparative Analysis of Susceptibility and Resistance Pattern of Antimicrobials. Archives of Clinical Microbiology, 7:4. doi:10.4172/1989-8436.100054.

3. Sabir S., Anjum A. A., Ijaz T., Ali M. A., Khan M. R., Nawaz M. (2014). Isolation and antibiotic susceptibility of E. coli from urinary tract infections in a tertiary care hospital. Pakistan Journal of Medical Sciences, 30 (2):389–392.

4. Farshad S., Anvarinejad M., Tavana A. M., Ranjbar R., Japoni A., Zadegan R. M., Alborzi, A (2011), ‘Molecular epidemiology of Escherichia coli strains isolated from children with community acquired urinary tract infections. African Journal of Microbiology Research, 5 (26): 4476 – 4483.

5. Russo, T. A. & Johnson, J. R. (2009), Extraintestinal Pathogenic *Escherichia coli’*, In Alan, DTB & Lawrence, RS, Vaccines for Biodefense and Emerging and Neglected Diseases, Academic Press, London, pg. 939–961.

6. Johnson, J. R. & Stell, A. L. (2000). ‘Extended virulence genotypes of *Escherichia coli* strains from patients with urosepsis in relation to phylogeny and host compromise. Journal of Infectious Diseases, 181 (1): 261 – 272.

7. Belanger L., Garenaux A., Harel J., Boulianne M., Nadeau E. & Dozois, C. M.(2011). Escherichia colifrom animal reservoirs as a potential source of human extraintestinal pathogenic *E. coli*’, FEMS Immunology and Medical Microbiology, 62 (1): 1 – 10.

8. Kohler C. D. & Dobrindt U. (2011). ‘What defines extraintestinal pathogenic Escherichia coli? International Journal of Medical Microbiology, 301 (8), pp. 642 – 647.

9. Johnson J. R. & Russo T. A. (2002). ‘Extraintestinal pathogenic *Escherichia coli*: ‘The other bad *E. coli*’, Journal of Laboratory and Clinical Medicine, 139 (3): 155 – 162.

10. Graziani C., Luzzi I., Corro M., Tomei F., Parisi G., Giufre M., Morabito S., Caprioli A. & Cerquetti M. (2009). ‘Phylogenetic background and virulence genotype of ciprofloxacin-susceptible and ciprofloxacin-resistant *Escherichia coli* strains of human and avian origin. Journal of Infectious Diseases, 199 (8):1209 – 1217.

11. Johnson T.J., Wannemuehler Y., Johnson S. J., Stell A. L., Doetkott C., Johnson J.R., Kim K. S., Spanjaard L. & Nolan L. K. (2008). ‘Comparison of extraintestinal pathogenic *Escherichia coli* strains from human and avian sources reveals a mixed subset representing potential zoonotic pathogens. Applied and Environmental Microbiology, 74 (22): 7043 – 7050.

12. Ramos N. L., Saayman M. L., Chapman T. A., Tucker J. R., Smith H. V., Faoagali J., et al.(2010). Genetic relatedness and virulence gene profiles of *Escherichia coli strains* isolated form septiaemic and uroseptic patients. European Journal of Clinical Microbiology & Infectious Diseases, 29:15–23.

13. Matuszkiewicz-Rowińska J., Małyszko J., Wieliczko M. (2015). Urinary tract infections in pregnancy: old and new unresolved diagnostic and therapeutic problems, Archives of Medical Science, 11 (1): 67–77.

14. McNally A., Alhashash F., Collins M., Alqasim A., Paszckiewicz K., Weston V. and Diggle M. (2013) Genomic analysis of extra-intestinal pathogenic *Escherichia coli* urosepsis. Clinical Microbiology and Infection, 19:E328–E334.

15. Salipante S. J., Roach D. J., Kitzman, Snyder M. W., Stackhouse B., Butler-Wu S. M., Lee C., Cookson B. T., Shendure J. (2014). Large-scale genomic sequencing of extraintestinal pathogenic *Escherichia coli* strains. Genome Research, 25:119–128.

16. World Health Organization Media Release: WHO’s first global report on antibiotic resistance reveals serious, worldwide threat to public health, Source: http://www.who.int/mediacentre/news/releases/2014/amr-report/en, accessed: 6/25/2014

17. Gilbert N. M., O’Brien V. P., Scott Hultgren, Macones G., Lewis W. G., Lewis A. L. (2013). Urinary Tract Infection as a Preventable Cause of Pregnancy Complications: Opportunities, Challenges, and a Global Call to Action. Global Advances in Health and Medicine, 2 (5): 59–69.

18. Mukherjee M., Koley S., Mukherjee S., Basu S., Ghosh B., Chakraborty S. (2015). Phylogenetic background of *E. coli* isolated from asymptomatic pregnant women from Kolkata, The Journal of Infection in Developing Countries, 9 (7): 720–724.

19. Al-Mayahie S.M. (2013).Phenotypic and genotypic comparison of ESBL production by vaginal *Escherichia coli* isolates from pregnant and non-pregnant women. Annals of Clinical Microbiology and Antimicrobials. 25; 12:7. doi: 10.1186/1476-0711-12-7.

20. Alemu A., Moges F., Shiferaw Y., Tafess K., Kassu A., Anagaw B., and Agegn A. (2012). Bacterial profile and drug susceptibility pattern of urinary tract infection in pregnant women at University of Gondar Teaching Hospital, Northwest Ethiopia. BMC Research Notes, 5:197.

21. Onoh R. C., Umeora O. U. J., Egwuatu V. E., Ezeonu P. O., & Onoh T. J. P. (2013). Antibiotic sensitivity pattern of Uropathogens from pregnant women with urinary tract infection in Abakaliki, Nigeria. Infection and Drug Resistance, 6; 225–233.

22. Newman M. J., Frimpong E., Donkor E. S., Opintan J. A., Asamoah-Adu A. (2011) Resistance to antimicrobial drugs in Ghana. Infection and Drug Resistance, 4: 215 – 220.

23. Donkor E. S., Akumwena A., Amoo K. P, Owolabi M. O., Aspelund T, Gudnason V. (2016). Post-stroke bacteriuria among stroke patients attending a physiotherapy clinic in Ghana: a cross-sectional study. Therapeutics and Clinical Risk Management, 12:457–462.

24. Donkor E. S., Osei J. A., Anim-Baidoo I., Darkwah S. (2007). Risk of Asymptomatic Bacteriuria among People with Sickle Cell Disease in Accra, Ghana. Diseases, 5 (4)

25. Ghana Statistical Service. 2012. 2010 Population and Housing Census. Summary Report of Final Results. GSS, Accra.

26. GhanaWeb, 2018, Source: https://www.ghanaweb.com/GhanaHomePage/general/statistics.php

27. Japheth A Opintan, Mercy J Newman, Reuben E Arhin, Eric S Donkor, Martha Gyansa-Lutterodt, William Mills-Pappoe (2015). Laboratory-based nationwide surveillance of antimicrobial resistance in Ghana. Infection and Drug Resistance, 8:379–389.

28. Cheesbrough M. (2007). District laboratory practice in tropical countries. Cambridge University Press. Part 2 pg. 71 – 140.

29. CLSI. Performance Stardards for Antimicrobial Disk Diffusion Tests (2015). Approved Stardard - 12^th^ Ed. CLSI documents MO2 – 12, Wayne, PA: Clinical and Laboratory Stardards Institutes.

30. Johnson, J. R., et al. (2006). Experimental mouse lethality of *Escherichia coli* isolates, in relation to accessory traits, phylogenetic group, and ecological source. Journal of Infectious Diseases, 194:1141–1150.

31. Lefort A., Panhard X., Clermont O et al. (2011). Host factors and portal of entry outweigh bacterial determinants to predict the severity of *Escherichia coli* bacteraemia. Journal of Clinical Microbiology, 49:777–783.

32. Maynard C, Bekal S., Sanschagrin F., Levesque R. C., Brousseau R., Masson L., Lariviere S., Harel J. (2004) Heterogeneity among virulence and antimicrobial resistance gene profiles of Extraintestinal *Escherichia coli* isolates of animal and human origin. Journal of Clinical Microbiology. 42 (12): 5444 – 5452.

33. Saenz Y, Brinas L, Dominguez E, Ruiz J, Zarazaga M, Vila J, Torres C. (2004) Mechanisms of resistance in multiple-antibiotic-resistant *Escherichia coli* strains of human, animal, and food origins. Antimicrobial Agents and Chemotherapy. 48 (10): 3996 – 4001.

34. Johnson JR, Russo TA, Tarr PI, Carlino U, Bilge SS, Vary JC. Jr, Stell AL. (2000) Molecular epidemiological and phylogenetic associations of two novel putative virulence genes, *iha* and *iroN*(*E. coli*), among *Escherichia coli* isolates from patients with urosepsis. Infect Immun. 68(5):3040 – 30407.

35. Johnson JR., Gajewski A, Lesse AJ, Russo TA. (2003). Extraintestinal Pathogenic *Escherichia coli* as a Cause of Invasive Non urinary Infections. Journal of Clinical Microbiology, 41(12), 5798–580.

36. Skyberg JA, Horne SM, Giddings CW, Wooley RE, Gibbs PS, Nolan LK. (2003). Characterizing avian *Escherichia coli* isolates with multiplex polymerase chain reaction. Avian Dis. 47: 1441–1447.

37. Bauer RJ, Zhang L, Foxman B, Siitonen A, Jantunen ME, Saxen H, Marrs CF. (2002) Molecular Epidemiology of 3 Putative Virulence Genes for *Escherichia coli* Urinary Tract Infection–*usp*, *iha*, and *iroN*E. The Journal of Infectious Diseases. 185 (10):1521–1524.

38. Paixão AC, Ferreira AC, Fontes M, Themudo P, Albuquerque T, Soares MC, Fevereiro M, Martins L, Corrêa de Sá MI. (2016) Detection of virulence-associated genes in pathogenic and commensal avian *Escherichia coli* isolates. Poultry Science. 95 (7):1646–1652.

39. Guiral E, Bosch J, Vila J, Soto S M. Prevalence of *Escherichia coli* among samples collected from the genital tract in pregnant and non pregnant women: relationship with virulence. FEMS Microbiol Lett. 2011; 314: 170–173.

40. Watt S, Lanotte P, Mereghetti L, Moulin-schouleur M, Picard B, Quentin R. *Escherichia coli* Strains from Pregnant Women and Neonates: Intraspecies Genetic Distribution and Prevalence of Virulence Factors. J. Clin. Microbiol. 2003; 41:1929–1935.

41. Sáez-López E, Guiral E, Fernández-Orth D, Villanueva S, Goncé A, López M, et al. (2016) Vaginal versus Obstetric Infection *Escherichia coli* Isolates among Pregnant Women: Antimicrobial Resistance and Genetic Virulence Profile. PLoS ONE 11(1):e0146531.doi:10.1371/journal.pone.0146531.

42. Boye A, Siakwa PM, Boampong JN, Koffuor GA, Ephraim RKD, Amoateng P, Obodai G, Penu D. (2012). Asymptomatic urinary tract infections in pregnant women attending antenatal clinic in Cape Coast, Ghana. Journal of Medical Research, 1 (6): 74 – 83.

43. Okonko IO. Donbraye-Emmanuel OB, Ijandipe LA, Ogun AA, Adedeji AO, Udeze AO. (2009). Antibiotics Sensitivity and Resistance Patterns of Uropathogens to Nitrofurantoin and Nalidixic Acid in Pregnant Women with Urinary Tract Infections in Ibadan, Nigeria. Middle-East Journal of Scientific Research. 4 (2): 105–109.

44. Turay AA, Eke SO, Oleghe PO, Ozekhome MC. (2014.)The prevalence of urinary tract infections among pregnant women attending antenatal clinic at Ujoelen primary health care centre, Ekpoma, Edo state, Nigeria. IJBAIR, 3 (1): 86 – 94.

45. Turpin C. A., Minkah B., Danso K. A., Frimpong E. H. (2007). Asymptomatic bacteriuria in pregnant women attending antenatal clinic at Komfo Anokye teaching hospital, Kumasi. Ghana Medical Journal. 41(1):26–9.

46. Lumbiganon P., Villar J., Laopaiboon M., Widmer M., Thinkhamrop J., Carroli G., Duc Vy N., Mignini L., Festin M., Prasertcharoensuk W., Limpongsanurak S., Liabsuetrakul T., Sirivatanapa P. (2009). One-day compared with 7-day nitrofurantoin for asymptomatic bacteriuria in pregnancy: a randomized controlled trial. Obstetrics and Gynecology. 113 (2 Pt 1):339–45.

47. Tadesse E., Teshome M., Merid Y., Kibret B., Shimelis T. (2014). Asymptomatic urinary tract infection among pregnant women attending the antenatal clinic of Hawassa Referral Hospital, Southern Ethiopia. BMC Research Notes, (7) 155.

48. Tazebew D., Getenet B., Selabat M., Wondewosen T. (2012). Urinary bacterial profile and antibiotic susceptibility pattern among pregnant women in North West Ethiopia. Ethiopian Journal of Health Science, 22 (2), 121–128.

49. Hamdan Z. H., Abdel Haliem M Ziad, Salah K. Ali, Ishag Adam, (2011). Epidemiology of urinary tract infections and antibiotics sensitivity among pregnant women at Khartoum North Hospital. Annals of Clinical Microbiology and Antimicrobial, 10:2.

50. Kawser P., Afroza M., Arzumath A. B., Monowara B, (2011). Prevalence Of Urinary Tract Infection During Pregnancy. Journal of Dhaka National Medical College Hospital. 17 (02): 8–12.

51. Akobi O. A., Inyinbor H. E., Akobi E. C., Emumwen E. G., Ogedengbe S.O., Uzoigwe E. O., Agbayani R. O., Emumwen E. F., Okorie I. E. (2014). Incidence of Urinary Tract Infection among Pregnant Women Attending Antenatal Clinic at Federal Medical Centre, Bida, Niger-State, North Central Nigeria. American Journal of Infectious Diseases and Microbiology, 2 (2): 34–38.

52. Sheerin N. S. (2015). Urinary tract infection. (Obstruction and Infection), Medicine, 43 (8): 435 – 439.

53. Emiru T., Beyene G., Tsegaye W. and Melaku S. (2013). Associated risk factors of urinary tract infection among pregnant women at Felege Hiwot Referral Hospital, Bahir Dar, North West Ethiopia. BMC Research Notes, 6:292.

54. Enayat K., Fariba F, Bahram N. (2008): Asymptomatic bacteriuria among pregnant women referred to outpatient clinics in Sanandaj, Iran. International Brazilian Journal of Urology 34:699–707.

55. Amiri FN, Rooshan MH, & Ahmady MH. (2009): Hygiene practices and sexual activity associated with urinary tract infection in pregnant women. East Mediterranean Health, 15:104–110

56. Ferede G, Yismaw G, Wondimeneh Y, Sisay Z. (2012). The Prevalence and Antimicrobial Susceptibility pattern of Bacterial Uropathogens Isolated from pregnant women. European Journal of Experimental Biology, 2 (5):1497‒1502.

57. Al-Haddad AM. (2005). Urinary tract infection among pregnant women in Al-Mukalla district, Yemen. East Mediterranean Health Journal. 11(3):505–510.

58. Sibi G, Kumari P. & Neema K., (2014). Antibiotic sensitivity pattern from pregnant women with urinary tract infection in Bangalore, India. Asian Pacific Journal of Tropical Medicine. 7 (Suppl 1): S116–S120.

59. Nandy P, Thakur AR, Ray CS. (2007). Characterization of bacterial strains isolated through microbial profiling of urine samples. OnLine Journal of Biological Sciences. 7, 44–51.

60. Feglo P. (2007). Antimicrobial sensitivity patterns of urine isolates at the Komfo-Anokye Teaching Hospital, (KATH) Kumasi, 2000-2005. Ghana Journal of Allied Health Sciences, 1 (2.): 43–49.

61. Newman MJ, Frimpong E, Asamoah-Adu A., Donkor ES. Resistance to antimicrobial drugs in Ghana. The Ghanaian-Dutch Collaboration for Health Research and Development, 2006; pp. 1–19, Source: https://www.researchgate.net/profile/Japheth_Opintan/publication/221760926_Resistance_to_antimicrobial_drugs_in_Ghana/links/09e41502445a36821a000000/Resistance-to-antimicrobial-drugs-in-Ghana.pdf

62. Johnson JR, Delavari P, O’Bryan TT, Smith KE, Tatini, S, Contamination of retail foods, particularly Turkey, from community markets (Minnesota, 1999 - 2000) with antimicrobial-resistant and extraintestinal pathogenic *Escherichia coli*, Foodborne Pathogens and Disease; 2005, 2(1): 38 – 49.

63. Johnson JR, McCabe JS, White DG, Johnston B, Kuskowski MA, McDermott P. (2009) Molecular analysis of *Escherichia coli* from retail meats (2002-2004) from the United States national antimicrobial resistance monitoring system, Clinical Infectious Diseases. 49 (2): 195 – 201.

64. Motayo BO, Ogiogwa IJ, Okerentugba PO, Innocent-Adiele HC, Nwanze JC, Onoh CC, et. al. (2012) Antimicrobial Resistance Profile of Extra-intestinal Escherichia coli Infections in a South Western Nigerian City. J. Microbiol Res. 2:141–144.

65. Mshana SE, Matee M, Rweyemamu M. (2013) Antimicrobial resistance in human and animal pathogens in Zambia, Democratic Republic of Congo, Mozambique and Tanzania: an urgent need of a sustainable surveillance system. Ann. Clin. Microbiol. Antimicrob. 12:28.

66. Ramos NL, Sekikubo M, Dzung DTN, Kosnopfel C, Kironde F, Mirembe F, Braunera A. (2012) Uropathogenic *Escherichia coli* Isolates from Pregnant Women in Different Countries. Journal of Clinical Microbiology. 50 (11):3569–3574

